# Determination of nucleotide-nucleotide and nucleotide-amino acid binding interactions from all-atom potential-of-mean-force calculations

**DOI:** 10.1101/2025.10.27.684844

**Authors:** Alejandro Feito, Eduardo Pedraza, Estefania Cuesta, Alejandro Castro, Ignacio Sanchez-Burgos, Antonio Rey, Rosana Collepardo-Guevara, Andrés R. Tejedor, Jorge R. Espinosa

## Abstract

Biomolecular condensates emerge from multivalent interactions between proteins and nucleic acids and are frequently modeled using coarse-grained molecular dynamics simulations. The parametrization of these models critically depends on atomistic data describing the underlying molecular interactions. In this work, we employ all-atom molecular dynamics simulations and potential-of-mean-force (PMF) calculations to investigate the interaction landscape between RNA nucleotides and protein amino acids. We begin by characterizing nucleotide–nucleotide binding modes through canonical base-pairing analysis, observing notable agreement in the predictions from both AMBER03ws and CHARMM36 force fields. Further rationalization of different nucleotide–nucleotide interaction modes involves the calculation of PMFs for ribose–ribose, phosphate–phosphate, and RNA tertiary interactions such as G-quadruplex formation. We also examine the effect of salt concentration on these interactions, finding a reduction on electrostatic self-repulsion for phosphate–phosphate binding upon increasing the ionic strength. Extending our analysis to amino acids, we first benchmark the performance of both AMBER03ws and a99SB-disp force fields for describing pairwise amino acid interactions, and then we evaluate different nucleotide–amino acid binding profiles. Our findings reveal a subset of amino acids—Lys and Arg (positively charged), Asp and Glu (negatively charged), and Gln, Ser, and Asn (polar residues)—that consistently engage with the nitrogenous bases of different nucleotides. Such binding is primarily mediated by hydrogen bonding and, in some cases, cation–*π* interactions. Furthermore, we identify strong *π*–*π* stacking interactions with aromatic residues and phosphate–Arg contacts as key contributors to condensate cohesion in RNA-protein condensates. Our comprehensive analysis provides a detailed library of nucleotide–amino acid interactions, offering quantitative insights to inform coarse-grained model parametrization and deepening our understanding of condensate self-assembly, nucleic acid recognition, and phase-separation regulation at submolecular scale.

## INTRODUCTION

Intracellular compartmentalization is a fundamental process that enables cells to organize their components in space and time to perform diverse biological functions. This biomolecular organization relies on the coordinated dynamics of both membrane-bound and membraneless organelles, the latter also referred to as biomolecular condensates. These condensates are thought to form through the spontaneous self-assembly of protein and nucleic acid solutions^1–3^—a mechanism that has attracted considerable attention across disciplines due to its broad implications for health and disease. Condensate formation allows cells to establish highly concentrated, yet transient, biomolecular assemblies—such as Cajal bodies^4^, stress granules^5^, P bodies^6^, and nucleoli^7^—that facilitate essential processes including biomolecular compartmentalization^8^, regulation of gene expression^9^, cellular stress response^10^, and signal transduction^11^. Given that many biomolecular condensates are primarily composed of proteins and nucleic acids (in the form of RNAs or DNAs), it is crucial to highlight the central role of their interactions in driving self-assembly and organization through phase-separation^12–14^. Importantly, transitions from dynamic, functional biomolecular condensates to irreversible, solid-like aggregates are mediated by the local strengthening of inter-protein ^15–17^ and RNA-RNA interactions^18,19^. Some of these transitions have been associated with the development of multiple neurodegenerative disorders, including amyotrophic lateral sclerosis (ALS) or frontotemporal dementia (FTD)^20,21^ among others^22^, leading to aberrant condensate formation^23^, fibril aggregation^24^, and ultimately, cytotoxicity^25^.

In the complex network of molecular interactions found in biomolecular condensates, electrostatic interactions play a central role in regulating the condensation of many different biomolecules^26,27^. In this context, RNA has been identified as a primary electrostatically driven condensate modulator^26,28–31^, capable of binding protein RNA-recognition motifs rich in positively charged^27,32–34^ and polar^35^ amino acids. More importantly, recent *in vitro* assays suggest that RNA alone is able to undergo phase-separation^18,19,36^, favored by low concentrations of magnesium ions that screen phosphate–phosphate repulsion. Furthermore, tertiary interactions such as those in G-quadruplex (G4) structures—non-canonical DNA/RNA formations composed of two or more guanine tetrads connected by Hoogsteen hydrogen bonds in a square planar arrangement^18,37,38^—have been associated with RNA aberrant aggregation leading to kinetically trapped states^19^. The precise interactions underlying the formation of RNA condensates and the role of salt concentration have not yet been fully explored^19,39^, although it is well established that *π*–*π* stacking^40^, hydrogen bonding^41^, and cation–*π* interactions^42,43^ modulate RNA and RNA–protein associations, thereby regulating key physicochemical properties of condensates such as viscosity and surface tension^44–46^.

In the last decade, molecular dynamics (MD) simulations have become a fundamental tool for studying condensate formation, providing mechanistic insights into their phase behavior, time-dependent material properties, and the key residues governing these processes^47–51^. In particular, residue-resolution coarse-grained (CG) models such as the CALVADOS2^52^, HPS-Urry^53^, or the Mpipi-Recharged^54–56^ have significantly advanced our ability to determine the physicochemical properties of biomolecular condensates in a precise and computationally efficient manner^52,57^. The pioneering HPS model and its subsequent parameterizations were mainly developed to reproduce experimental radii of gyration and to qualitatively capture the phase behavior of different proteins^53,58,59^. However, the focus of newly designed models has shifted towards bottomup approaches that leverage atomistic simulations^54,60^, bioinformatics^61^, or machine-learning-based parameterizations^52,62–65^. In particular, potential-of-mean-force (PMF) calculations have gained prominence for evaluating the free energy of binding in various biological systems, including peptides^50,66,67^, amino acids^54,60,61,68–70^, and RNA–protein interactions^54^. These approaches provide submolecular insights into the mechanisms and interactions underlying condensate formation or aberrant liquid-to-solid transitions^47,68,71^.

In this work, we perform a systematic characterization of RNA nucleotide pairs and nucleotide–amino acid interactions by means of all-atom PMF calculations with explicit solvent and ions. We first benchmark representative atomistic force fields—AMBER03ws^72^ and CHARMM36/TIP4P2005^73,74^—for describing canonical base-pairing and verify that both reproduce the expected hydrogen-bonding hierarchy (CG *>* AU *>* GU). We analyze potential interaction modes of nucleotides by calculating their PMF profiles at varying relative orientations. In particular, we estimate binding interaction strengths for base–base, phosphate–phosphate, and ribose–ribose interactions. Moreover, RNA tertiary interactions involving G-quadruplexes are also included in our PMF calculation scheme to further characterize more complex RNA structures. We then explore how specific nucleotide–nucleotide affinities depend on the concentration of monovalent salt (0–3 M NaCl), demonstrating that while base–base and ribose–ribose interactions mostly remain unaltered, high salt concentrations effectively screen phosphate–phosphate repulsion. We also extend our calculations to encompass interactions between the 20 natural amino acids and the four RNA nucleotides, covering all possible combinations. To this end, we first validate the AMBER03ws/TIP4P2005 force field^72,74^ against the a99SB-*disp*/TIP4P-*disp*^75^ model to ensure accurate reproduction of pairwise amino acid interactions. We then determine the binding of amino acids to nucleotides, focusing on those relevant for biomolecular condensate formation, including interactions between phosphate groups and charged residues, and *π*–*π* stacking of aromatic residues with the base and ribose. Overall, our results advance the understanding of nucleotide–amino acid binding, providing an extensive library that serves as a foundation for the future development of improved coarse-grained models of protein and nucleic acid condensates, and for rationalizing experimental *in vitro* and *in vivo* observations at the molecular level.

## I. METHODOLOGY

The primary focus of this study is to rationalize the molecular interactions between the residues and nucleotides involved in biomolecular condensation. To this end, atomistic PMF calculations were performed using the GROMACS simulation package (version 2023)^76^. The molecular interactions of amino acids and nucleotides—including nucleotide–nucleotide, amino acid–amino acid, and nucleotide–amino acid— were modeled using the AMBER03ws/TIP4P2005 force field^72,74^, and validated by contrasting with the CHARMM36/TIP4P2005^73,74^ and a99SB-*disp*/TIP4P-*disp*^75^ models for the nucleotide–nucleotide and amino acid–amino acid interactions, respectively. Note that we refer to the water model in a99SB-*disp* as TIP4P-*disp*, given that a99SB-*disp* is a co-parameterization of the AMBER99SB*-ILDN-Q^77^ protein force field together with the TIP4P-D water model^78^, which includes modifications to the C6 dispersion term of the water oxygen. All simulations were performed at 300 K and physiological salt concentration (150 mM NaCl). Additionally, the effect of salt concentration was explored in the range 0–3 M NaCl for nucleotide–nucleotide and amino acid–amino acid interactions. To build the systems, the phosphate group of each nucleotide was capped with a methyl group, while the N- and C-terminal ends of each amino acid were capped with acetyl and N-methyl groups, respectively.

For nucleotide–nucleotide PMFs, the nucleotide pair was pulled along a dissociation coordinate defined by the distance between the centers of mass (COMs) of the interacting structural regions: the nitrogenous base, the ribose, or the phosphate group. Thus, nucleotide pairs were oriented to face each other via the nitrogenous base, ribose, or phosphate, with the COM of the corresponding region serving as the reference in the initial configuration. For nucleotide–amino acid and amino acid–amino acid PMFs, the reaction coordinate was defined as the distance between the COMs of the most relevant regions of both molecules (e.g., bases, riboses, side chains, or aromatic rings). In nucleotide–amino acid PMFs, molecules were positioned such that the base faced the side chain of the amino acid. For amino acid–amino acid PMFs, the pairs were oriented to face through their side chains to enable potential binding. In all cases, the residue pairs were placed in a cubic box of dimensions 3.4 × 4.4 × 10 nm containing approximately 5000 water molecules. Some water molecules were replaced by Na^+^ and Cl^−^ ions to achieve the desired salt concentration and electroneutrality. Energy minimization was performed with a force tolerance of 1000 kJ mol^−1^ nm^−1^, applying positional restraints of 8000 kJ mol^−1^ nm^−2^ to all heavy atoms of the interacting residues.

During production runs, temperature coupling was achieved using the velocity-rescale (v-rescale) thermostat with a relaxation constant *τ*_*T*_ = 1.0 ps, and pressure coupling was applied via the Parrinello–Rahman barostat with *τ*_*P*_ = 1.0 ps. Positional constraints of 1000 kJ mol^−1^ nm^−2^ were applied in directions perpendicular to the pulling axis. The COM distance between nucleotide pairs was controlled using a harmonic umbrella potential with a force constant of 10000 kJ mol^−1^ nm^−2^, while a lower constant (6000 kJ mol^−1^ nm^−2^) was used for amino acid–amino acid interactions. Bond constraints on hydrogen-containing bonds were enforced using the LINCS algorithm, allowing an integration time step of 2 fs. Periodic boundary conditions (PBC) were maintained in all three spatial directions. Short-range dispersive and electrostatic interactions were calculated using a cutoff of 0.9 nm, and long-range contributions were computed with the Particle-Mesh Ewald method^79^. Approximately 30 umbrella sampling windows were defined, spaced every 0.05 nm, covering the full COM distance range. Each window was simulated for 10 ns, and the Weighted Histogram Analysis Method (WHAM)^80^, as implemented in GROMACS, was used to reconstruct the free-energy profiles. The initial 2000 ps of each simulation were discarded to ensure equilibration. For RNA tertiary interactions involving G-quadruplexes, we employed a PDB structure as the starting configuration. All parameters and conditions were identical to those described above, except that the pulling was applied to one strand while the remaining three were treated as a rigid block. The system was solvated in a rectangular box of dimensions 7 × 9 × 10 nm containing approximately 20,000 water molecules. The reaction coordinate was defined as the COM distance between the pulled strand and the rest of the quadruplex, and sampling was performed using a harmonic umbrella potential with a force constant of 55000 kJ mol^−1^ nm^−2^.

The binding free energy for all PMF simulations was estimated as the global minimum of the resulting free-energy curve. We verified that this approach yields equivalent values to integrating the negative part of the PMF, as confirmed for base–base and amino acid–amino acid systems in Figs. S2 and S5. To test the robustness of the positional constraints, additional PMF calculations allowing free rotation of both residues were also performed (Fig. S1). The uncertainty of the PMF minimum was estimated using the Bayesian bootstrap method^81^. To account for local fluctuations, the final error was obtained by combining the Bayesian bootstrap errors of the minimum and of the adjacent points.

## RESULTS

### A. Characterization of the different binding modes in nucleotide-nucleotide interactions

We systematically performed atomistic PMF calculations for different configurations of nucleotide pairs. With this approach, we unraveled the various binding modes between nucleotides and how they are modulated by their relative orientation and salt concentration. We first computed the PMFs for canonical Watson–Crick base pairs (i.e., CG and AU) and for the non-canonical GU wobble pair, interacting through the base in specific configurations extracted from experimental PDB structures. In particular, we used the PDB codes 1RNA for AU, 259D for CG, and 1A4D for GU. All simulations were performed using the AMBER03ws force field^72,82^, which enables intrinsically disordered proteins to explore extended conformational states, reduces non-specific aggregation, and provides a realistic description of solvation and the structural properties of disordered chains. In addition, AMBER03ws has been shown to realistically capture RNA binding to TDP-43^83^ and recapitulate the free energy of binding between different types of nucleotides^54,61^. As a benchmark, we also calculated the PMF profiles for the canonical base pairs using the CHARMM36 force field^73^, which has been specifically developed and validated for proteins^84^, lipids^85^, and nucleic acids^86^.

In Fig. 1A, we show the PMF profiles of the canonical Watson–Crick pairs (CG, AU) and the non-canonical wobble pair (GU) obtained with both force fields. The results reproduce the expected trend for Watson–Crick base pairing, corresponding to three versus two hydrogen bonds for CG and AU, respectively. The CG pair exhibits the deepest free energy minimum, followed by AU, maintaining a ratio of 0.8 between them, which is reasonably close to the expected 2/3 value derived from the number of hydrogen bonds. Importantly, both force fields yield similar profiles (∼ 0.5 kcal/mol), although the PMFs from AMBER03ws consistently display slightly deeper minima. This behavior can be attributed to the stronger interactions captured by AMBER03ws, as the discrepancies between the two models become more pronounced with increasing numbers of hydrogen bonds between the nucleotide pairs. Additionally, the GU wobble pair shows a significantly lower binding strength, as expected, and exhibits a similar shape for both force fields. Crucially, both models reproduce the same relative energetic order (CG *>* AU *>* GU), demonstrating the robustness of our approach. In fact, when normalizing the absolute minima of the interaction profiles by the binding energy of CG for each model (Fig. 1B), the relative differences between force fields become negligible. In the following calculations, we therefore employ the AMBER03ws model unless otherwise specified.

**FIG. 1.**
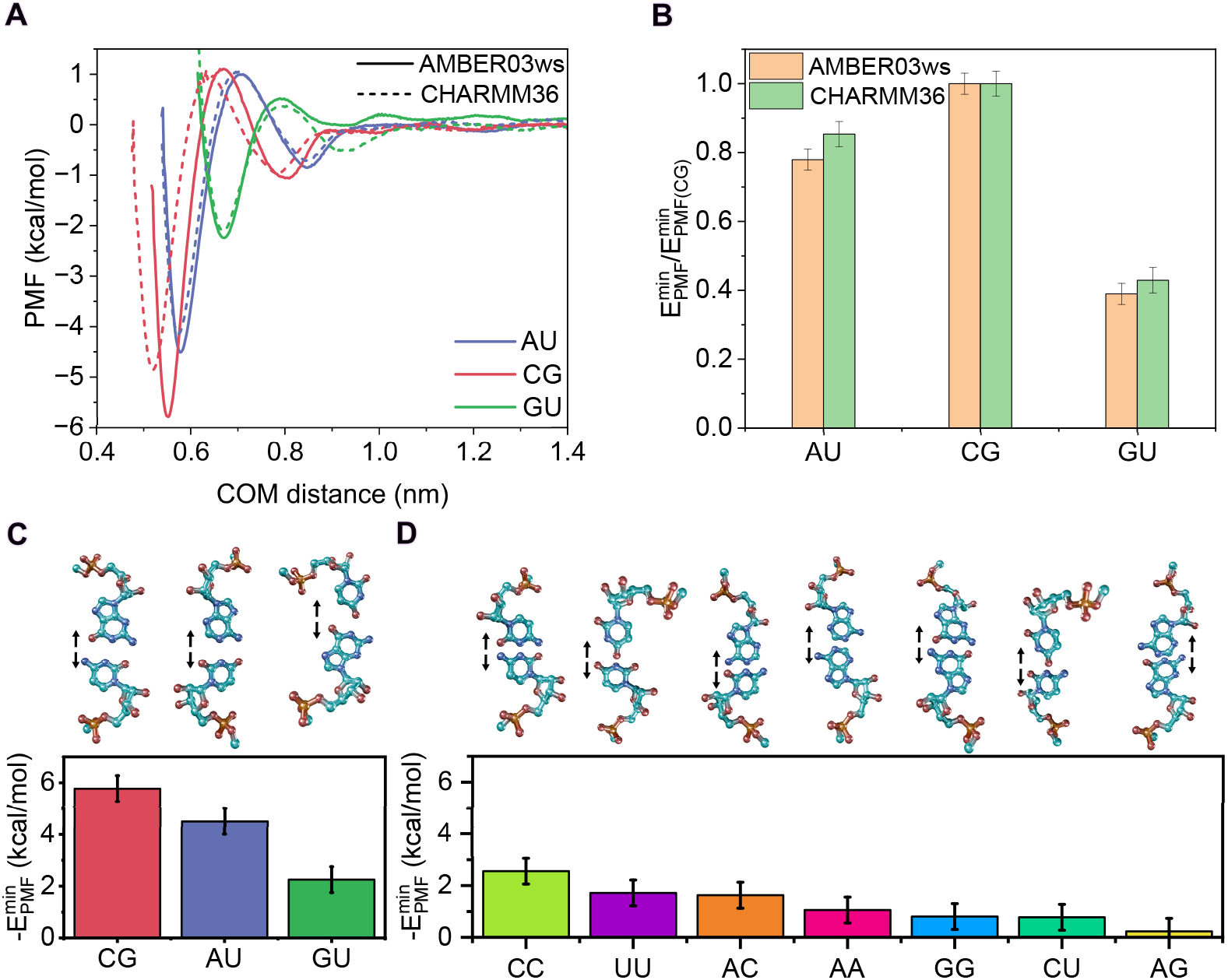
(**A**) PMF profiles as a function of the center-of-mass (COM) distance between the bases of the corresponding pair obtained with AMBER03ws (solid lines) and CHARMM36 (dotted lines) force fields for the canonical pairs adenine–uracil (AU, blue) and cytosine–guanine (CG, red), and the guanine–uracil ‘wobblé (GU, green) pair. (**B**) Free energies estimated from the minimum of the PMF profiles for canonical nucleotide pairs with AMBER03ws (orange bars) and CHARMM36 (green bars), and normalized by the value of the CG pair. Absolute minimum free energies of the PMF profiles for canonical and wobble (**C**) and non-canonical (**D**) nucleotide pairs. Above the energy plot we depict the nucleotide pair configurations used in the PMF calculations including arrows to indicate the dissociation direction with the atom color code: Light blue (carbon), dark blue (nitrogen), red (oxygen), and orange (phosphorus). All calculations were performed at T= 300 K, p= 1 bar, and 150 mM NaCl.

We next extend our free energy calculations to all remaining nucleotide combinations to estimate their interactions through the base (Fig. 1C–D). We report the binding free energies for the canonical and wobble pairs (AU, CG, and GU; Fig. 1C) as well as for the remaining non-canonical base pairs: CC, UU, AA, GG, AC, CU, and AG (Fig. 1D; see Fig. S2A in the Supplementary Material for the corresponding PMF profiles). For these non-canonical pairs, the configuration was set up by placing the residues in a mirror-image arrangement with the bases in the same plane, as described in Section I and illustrated in Fig. 1D. This configuration enables base–base interactions and facilitates hydrogen bond formation. As shown in Fig. 1D, the most energetically favorable pair is CC, consistent with cytosine’s ability to form hydrogen bonds^88,89^. The UU and AC pairs exhibit similar stability, slightly below that of CC, while the remaining pairs display interaction energies of the order of thermal fluctuations intrinsic to the system (∼ 0.5 kcal/mol; see Section I for further details on this estimation).

Complementary to our approach, some studies have used the C1’–C1’ (C1’ atom corresponds to the anomeric carbon of the ribose sugar, which is covalently bonded to the nucleobase) distance as a geometric descriptor correlated with nucleotide pairing type^90–92^, both for canonical Watson–Crick pairs and non-canonical pairs. Evaluating the C1^*′*^–C1^*′*^ distance provides insight into the stability of the pairs and can be readily computed from our simulations by modifying the reaction coordinate (see Fig. S2B in the Supplementary Material). We find that as the absolute interaction energy decreases (Fig. 1D), the C1^*′*^–C1^*′*^ distances tend to increase, indicating reduced stability for non-canonical base pairs relative to Watson–Crick pairs (see Fig. S2B), consistent with experimental observations^93,94^. Some authors have also proposed calculating the binding free energy from PMFs by integrating the negative part of the profile rather than taking the global minimum^54,61,68^. In Fig. S3, we show the resulting binding energies obtained using this integration approach for each nucleotide pair. The overall trend remains consistent across all pairs, although the relative binding strengths of AA, GG, and CU are slightly increased due to local minima in their PMF profiles at intermediate distances (see Fig. S2A).

Base–base interactions are crucial for nucleotide binding, particularly for pairs capable of forming persistent hydrogen bonds (i.e., CG, AU, GU, CC). However, the energy landscape of nucleotide interactions is not solely determined by the nitrogenous bases; interactions between other constituent groups, such as ribose and phosphate, can also exert a significant influence. Importantly, a systematic analysis of these interactions allows us to understand how the stability of nucleotide assemblies in various cellular organelles is affected by the different nucleotide groups. To this end, we performed PMF calculations evaluating ribose–ribose (Fig. 2A–B) and phosphate–phosphate (Fig. 2C–D) interactions, considering both *cis* and *trans* configurations (see snapshots in Fig. 2). In the *cis* configuration, the non-facing groups are arranged symmetrically, whereas in the *trans* configuration, the same groups are oriented to oppose each other (see snapshots in Figs. 2B,D). In general, ribose–ribose interaction profiles exhibit weak attractive energies for both *cis* and *trans* orientations (Fig. 2A), particularly when compared to canonical base pairing (Fig. 1). The *trans* configurations show slightly more attractive interactions than the *cis* configurations (Fig. 2B), except for the GU pair, likely due to the lower affinity between these nucleotides, which renders their ribose–ribose binding largely orientation-independent. Crucially, ribose–ribose interactions are largely insensitive to the specific nucleotide pair, as the magnitude of the well depth is comparable across the different studied cases. This observation suggests that ribose forms transient interactions rather than driving nucleotide association or RNA phaseseparation.

**FIG. 2.**
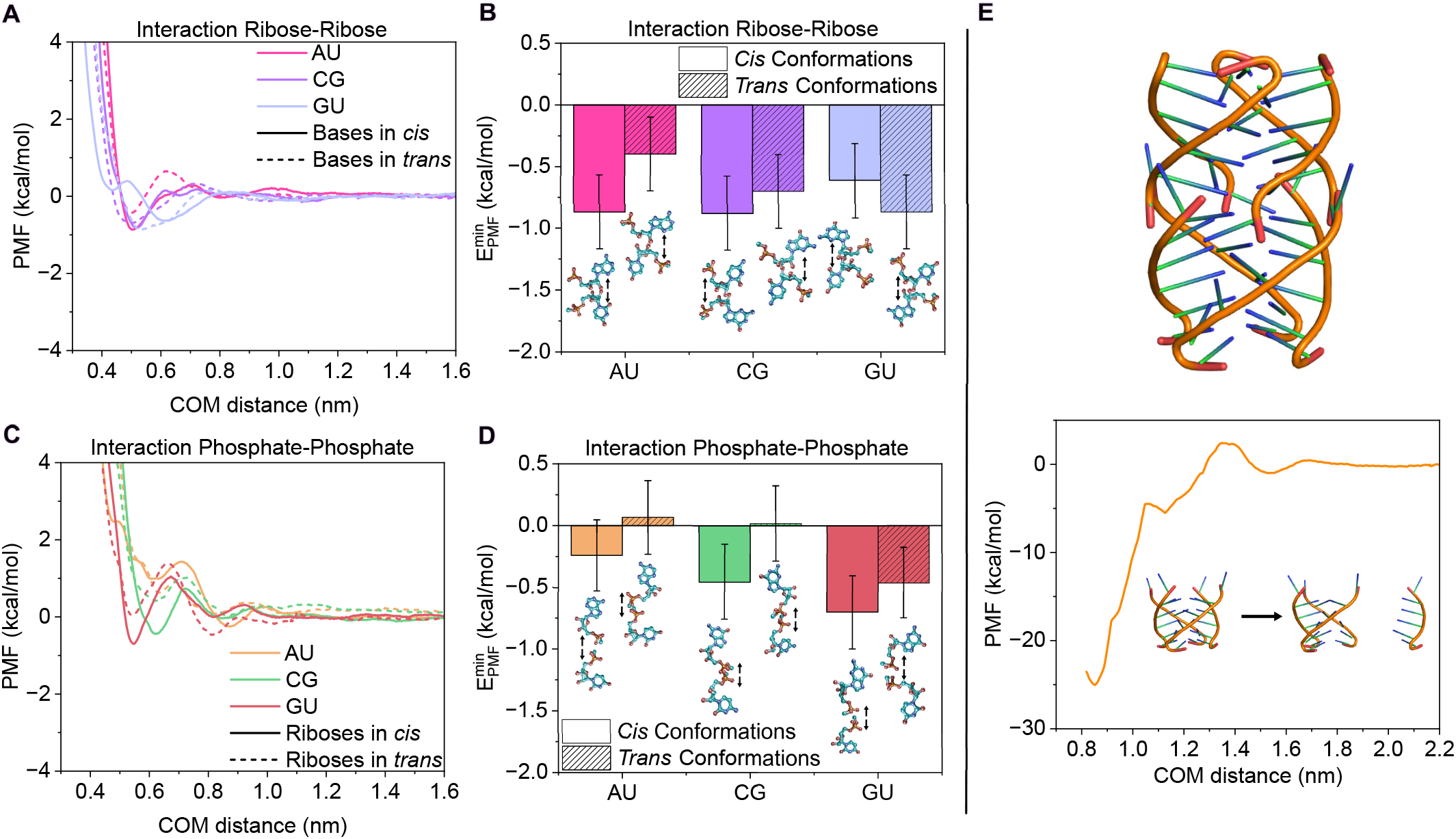
(**A**) PMF profiles as a function of the center-of-mass (COM) distance between the riboses of the canonical AU, CG and GU wobble pairs in *cis* (solid line) and *trans* conformation (dashed line). (**B**) Free energy interaction from the PMF minimum (panel **A**) for the configurations shown in the snapshots including arrows to indicate the dissociation direction. (**C**) PMF profiles as a function of the center-of-mass (COM) distance between the interacting phosphates of the canonical AU, CG and GU wobble pairs in *cis* (solid line) and *trans* conformation (dashed line). (**D**) Free energy interaction from the PMF minimum (panel **C**) for the configurations shown in the snapshots including arrows to indicate the dissociation direction. (**E**) Snapshot of the G-quadruplex structure from the experimental PDB code 4XK0^87^ (top) along with the PMF interaction profile as a function of the center-of-mass (COM) distance between the dissociated strand and the ternary cluster (bottom). All calculations were performed at T= 300 K, p= 1 bar, and 150 mM NaCl.

The phosphate–phosphate PMF profiles (Fig. 2C) provide insight into the electrostatic interactions arising from their negative charges. The curves reflect primarily repulsive interactions, with *cis* configurations exhibiting slightly larger negative energy values, in contrast to the *trans* configurations (Fig. 2D). It is important to note that the binding energy alone does not fully determine the interaction mode, as the shape of the PMF can also influence how the residues approach each other. In this context, the phosphate–phosphate PMF curves consistently display a repulsive barrier of ∼1–2 kcal/mol for all studied pairs, which hinders effective binding. Interestingly, the *trans* configuration of the GU pair shows an energy minimum of similar magnitude to that of the *cis* configuration, supporting the notion that guanine may promote relatively stable interactions between nucleotide pairs.

RNA and DNA secondary structures, governed by tertiary interactions such as G-quadruplexes, are essential for understanding the mechanisms underlying RNA liquid-like condensation versus solid-like pathological aggregation^19,95,96^. G-quadruplex (G4) structures form through Hoogsteen-type hydrogen bonds between the bases of four guanine nucleotides, creating a secondary structure that alters the local properties and geometry of the RNA (see Fig. 2E). The significance of this structure lies in its association with pathological processes, including cancer, neurodegenerative diseases, and viral replication, which also make G4s promising therapeutic targets^38,97,98^. To investigate the binding interaction strength of a G4 structure, we calculated its PMF profile using an experimental PDB structure (code 4XK0^87^; Fig. 2E). One strand forming the G4 was pulled while the remaining strands were restrained (see Section I for details). The resulting PMF curve shows a strong interaction between the strands, with a free energy minimum of ™25 kcal/mol at a COM distance of ≈ 0.85 nm. Each guanine is expected to form two hydrogen bonds via the N1–H donor group and the carbonyl oxygen at C6 as acceptor. Considering that each G4 strand contains six guanines, the binding energy associated with the dissociation of one guanine from each tetrad is approximately −4.2 kcal/mol. These values are alike to those found for canonical base pairs (Fig. 1A,C), consistent with the notion that the global stability of the G4 structure arises from the cooperative accumulation of multiple hydrogen bonds. For reference, intermolecular interactions within protein cross-*β*-sheets exhibit binding strengths of ∼15–40 kcal/mol in PMF simulations^49,66,67^, comparable to the energy estimated for the G-quadruplex, which can also contribute to kinetically arrested condensates as experimentally reported for G_4_C_2_ RNA repeats^18^.

Nucleotide interactions are influenced not only by their relative orientation but also by the salt concentration in the medium. Several studies have demonstrated the effect of salinity on the interaction profiles of amino acids^68,99^. Salt plays a crucial role in the stability and dynamics of biomolecular condensates by modulating multivalent interactions that can either promote or inhibit condensate formation in a context-dependent manner^48,54,100,101^. In particular, RNAs can undergo phase-separation at low concentrations of MgCl_2_ (∼ 10 mM), as the ions screen electrostatic self-repulsion and enable base-pairing^19,46,102,103^. To investigate the role of salt in modulating nucleotide-nucleotide interactions, we first benchmarked PMF profiles for different amino acid pairs at 0, 1.5, and 3 M NaCl, comparing our results with those of Krainer *et al*.^68^ (Fig. S4 in the Supplementary Material). Our configurations were designed to mimic the setup shown in Fig. 6 of the original work^68^. The resulting PMF curves quantitatively match those reported by Krainer *et al*., with only slight shifts in the distance scale for some pairs (e.g., Arg–Tyr, Arg–Glu). Importantly, when normalizing the global minimum energies by the highest value (Fig. S4G), the trends closely reproduce the original calculations, confirming that our force field implementation accurately captures the effect of salt concentration on residue-residue interactions.

We next extended the analysis to nucleotides, examining canonical base pairs at NaCl concentrations ranging from 0 to 3.00 M. In Fig. 3A–C, we show the free energy profiles for the canonical pairs at 0, 0.15, 0.50, 1.50, and 3.00 M, as indicated in the panels. For base–base interactions, the binding strength remains largely unchanged across all concentrations, with only a moderate decrease observed as salt increases from 0 to 3 M. A closer inspection of the energy minima (see insets in Figs. 2A–C) reveals that the minimum energy becomes less negative compared to the 0 M reference: by 0.8 kcal/mol for CG, 0.5 kcal/mol for AU, and 0.3 kcal/mol for GU. When normalizing the free energy minima by the value at 3 M NaCl (Fig. 3D), the slight progressive decrease in stability with increasing salt is highlighted, confirming that salt might only have a minor effect on hydrogen-bond-mediated base pairing.

**FIG. 3.**
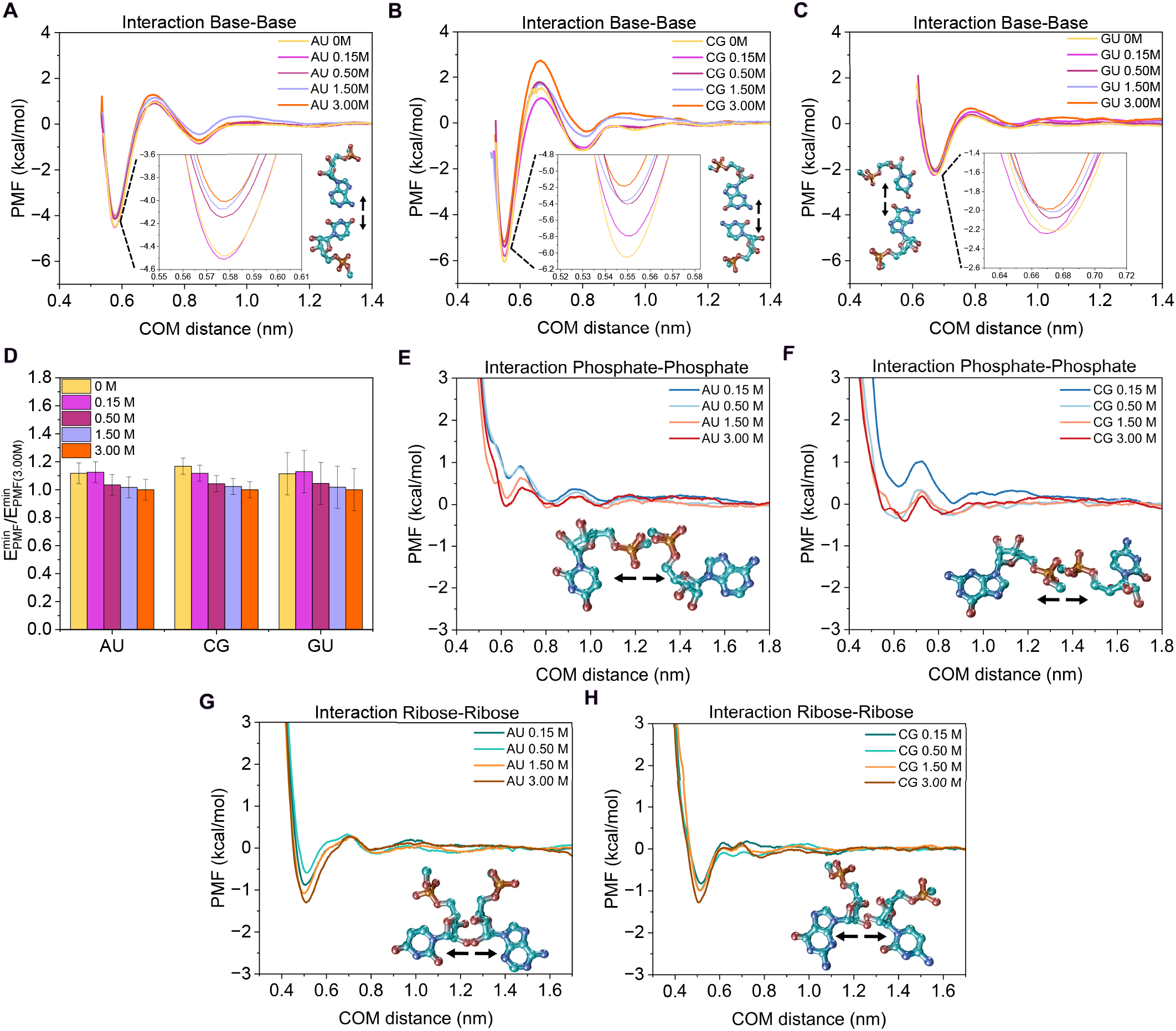
PMF profiles as a function of the center-of-mass (COM) distance between the bases obtained with AMBER03ws for the canonical nucleotide pairs AU (**A**), CG (**B**), and GU wobble (**C**), facing their nitrogenous bases and for several NaCl concentrations from 0-3 M. Insets show a zoom on the minima of the PMF profiles. (**D**) Free energies derived from the minima of the PMF profiles shown in panels **A**–**C** at various NaCl concentrations, normalized with respect to the minimum value of each pair at 3 M NaCl. PMF profiles as a function of the center-of-mass (COM) distance between the phosphates of the corresponding pair obtained at different salt concentrations concentrations for the canonical nucleotide pairs AU (**E**) and CG (**F**) oriented via their phosphate groups. PMF profiles as a function of the center-of-mass (COM) distance between the riboses for AU (**G**) and CG (**H**), and interacting through the riboses. Alongside the interaction profiles are the nucleotide pair configurations used in the PMF calculations including arrows to indicate the dissociation pathway.

Increasing the salt concentration is expected to primarily modulate electrostatic interactions, which in nucleotides are largely associated with the negatively charged phosphate groups. To explore this effect, we analyzed phosphate–phosphate interactions for the AU and CG pairs at NaCl concentrations ranging from 0.15 M to 3 M, using the *cis* configuration with phosphate groups aligned and facing each other (Figs. 2 and 3E–F).

As expected, the electrostatic repulsion is increasingly screened as salt concentration rises. In particular, the PMF for the CG pair (Fig. 3F) shows a notable reduction in repulsion at concentrations above 0.15 M NaCl. To further investigate this behavior, we constructed an alternative configuration in which the phosphate oxygens are directly aligned (see Fig. S5). In this setup, the AU pair exhibits a pronounced decay in the PMF with increasing NaCl, similar to the *cis* configuration of Fig. 3E. In contrast, the CG pair shows a nearly constant PMF minimum across all salt concentrations, although the curve smooths the energy barrier at intermediate distances (∼ 0.9 nm) and shifts the repulsive region to shorter distances (from ∼0.7 nm to ∼0.6 nm). Finally, we examined the influence of salt on ribose–ribose interactions for the canonical AU and CG pairs. Using a configuration where the ribose groups face each other symmetrically (Figs. 3G–H), we find that short-range interactions become slightly more attractive at higher salt concentrations. The resulting PMF profiles become significantly deeper than those observed for base–base interactions (Figs. 3A–C), with the free energy minimum increasing by approximately 0.5 kcal/mol, effectively doubling the well depth.

### B. Benchmark of amino acid pairwise interactions through different force fields

Proteins are major building blocks driving the formation of biomolecular condensates, making the calculation of amino acid–nucleotide interactions central to understanding phase-separation at the submolecular level. Before analyzing amino acid–nucleotide interactions, we first assessed whether amino acid interactions are similarly described by the AMBER03ws/TIP4P2005 force field, compared to a99SB-*disp*/TIP4P-*disp*, a widely used model for evaluating amino acid and peptide interactions^54,60,66,67,104^. To this end, we computed PMF profiles for five amino acid pairs abundant in phase-separating proteins: Ala–Ala, Arg–Arg, Arg–Asp, Arg–Tyr, and Tyr–Tyr, arranging the side chains to face each other (Figs. 4A–E; see Section I for details). The resulting PMFs show energetically consistent profiles between the two force fields and align well with the expected behavior^68,105^. Specifically, Arg–Tyr (cation–*π*^58^) and Tyr–Tyr (*π*–*π*^61,106^) exhibit the highest binding affinities. Differences between the force fields are also observed: AMBER03ws predicts deeper minima for Arg–Asp, Arg–Tyr, and Tyr–Tyr pairs compared to a99SB-*disp*, reflecting distinct capacities to capture intermolecular interactions between side chains^72,75^. Conversely, a99SB-*disp* favors attractive interactions between like-charged residues such as Arg–Arg, whereas AMBER03ws shows stronger affinity for oppositely charged pairs, such as Arg–Asp (Fig. 4C). Figure 4F summarizes these results, showing the PMF energy minima for each pair normalized by the Arg–Tyr interaction, the strongest residue–residue interaction stabilizing protein condensates^107^. Overall, while a99SB-*disp* yields slightly more attractive interactions for Ala–Ala and Arg–Arg, AMBER03ws predicts stronger binding for Arg–Asp and Tyr–Tyr, capturing trends relevant for protein condensate formation in a consistent manner.

**FIG. 4.**
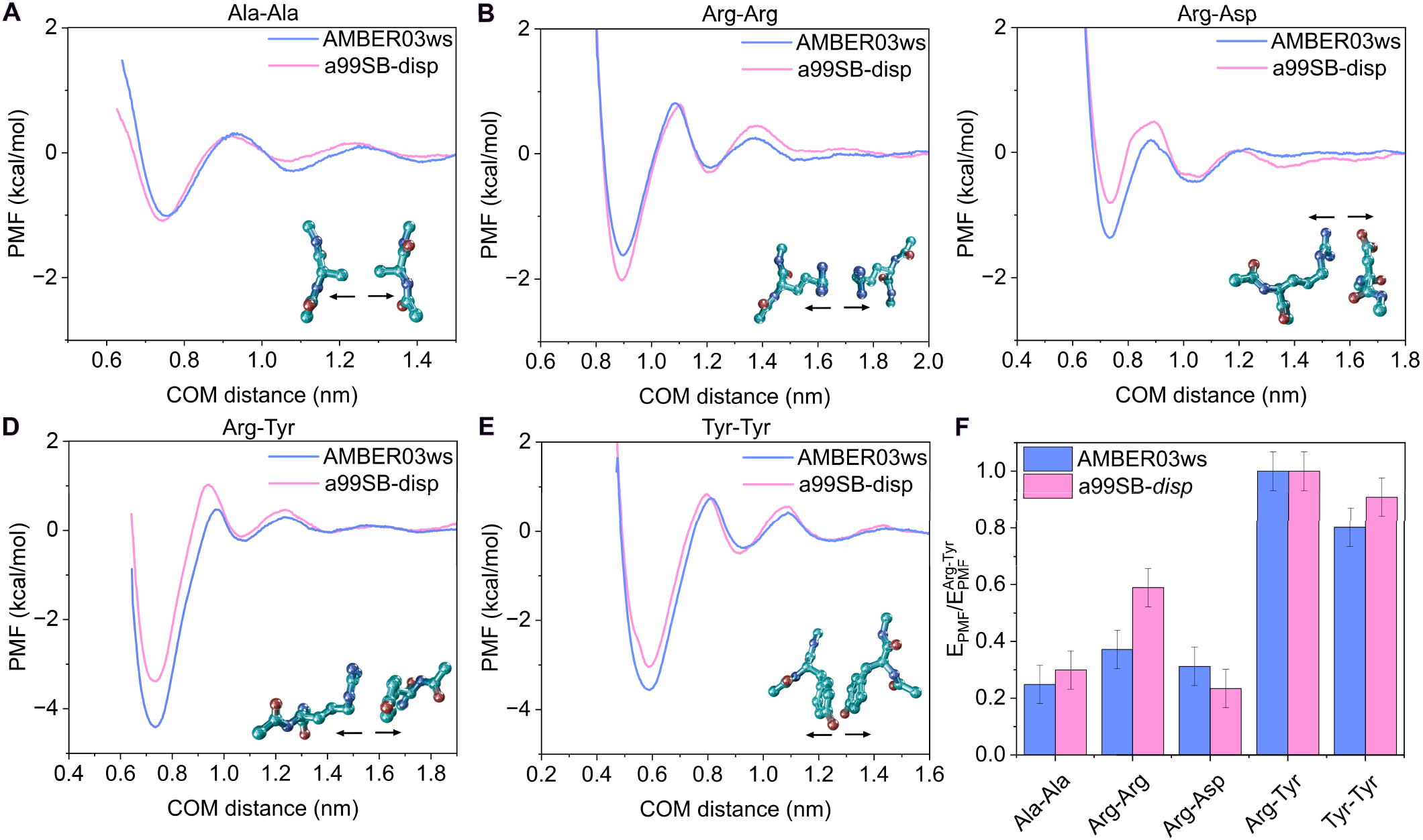
PMF profiles along the COM distance between the side chain of the corresponding interacting amino acids obtained with AMBER03ws (pink lines) and a99SB-*disp* (blue lines) for the amino acid pairs alanine–alanine (**A**), arginine–arginine (**B**), arginine–aspartic acid (**C**), arginine–tyrosine (**D**), and tyrosine–tyrosine (**E**), oriented to face through their side chains, at 300 K, 1 bar, and 150 mM NaCl. The amino acid pair configurations used in the calculations and the arrows to indicate the dissociation direction are shown next to the interaction profiles. (**F**) Free energy interaction from the minima of the PMF profiles for both models, normalized with respect to the binding energy of the arginine–tyrosine pair.

### C. Characterization of the nucleotide–amino acid interactions across distinct binding modes

RNA represents a fundamental modulator of biomolecular condensation by establishing heterotypic interactions with proteins that regulate condensate stability and material properties^26,28,49,108–111^. We therefore extend our PMF calculations to uncover the nucleotide–amino acid interactions underlying multivalent condensation. Specifically, we employ the AMBER03ws force field to characterize the interactions of the four RNA nucleotides (A, C, G, and U) with the twenty standard amino acids (Lys, Arg, Asp, Glu, His, Cys, Met, Phe, Tyr, Trp, Thr, Gln, Ser, Asn, Gly, Pro, Ala, Val, Leu, and Ile). As a first approach, and considering the large number of possible interaction modes, we systematically arranged each amino acid with its side chain oriented toward the nitrogenous base of the nucleotide (see Section I for technical details). The PMF interaction profiles and the corresponding free energy minima are shown in Figs. 5 and 6, together with representative snapshots of the employed configurations. As shown, nucleotide–amino acid interactions between the base and the side chain are generally weaker than canonical base–base interactions (Fig. 1), but comparable to amino acid–amino acid interactions under physiological conditions (see Fig. 4 and refs.^54,61,68^). Interestingly, seven amino acids (Lys, Arg, Asp, Glu, Gln, Ser, and Asn) exhibit significant interactions with most nucleotides, especially with C and G.

**FIG. 5.**
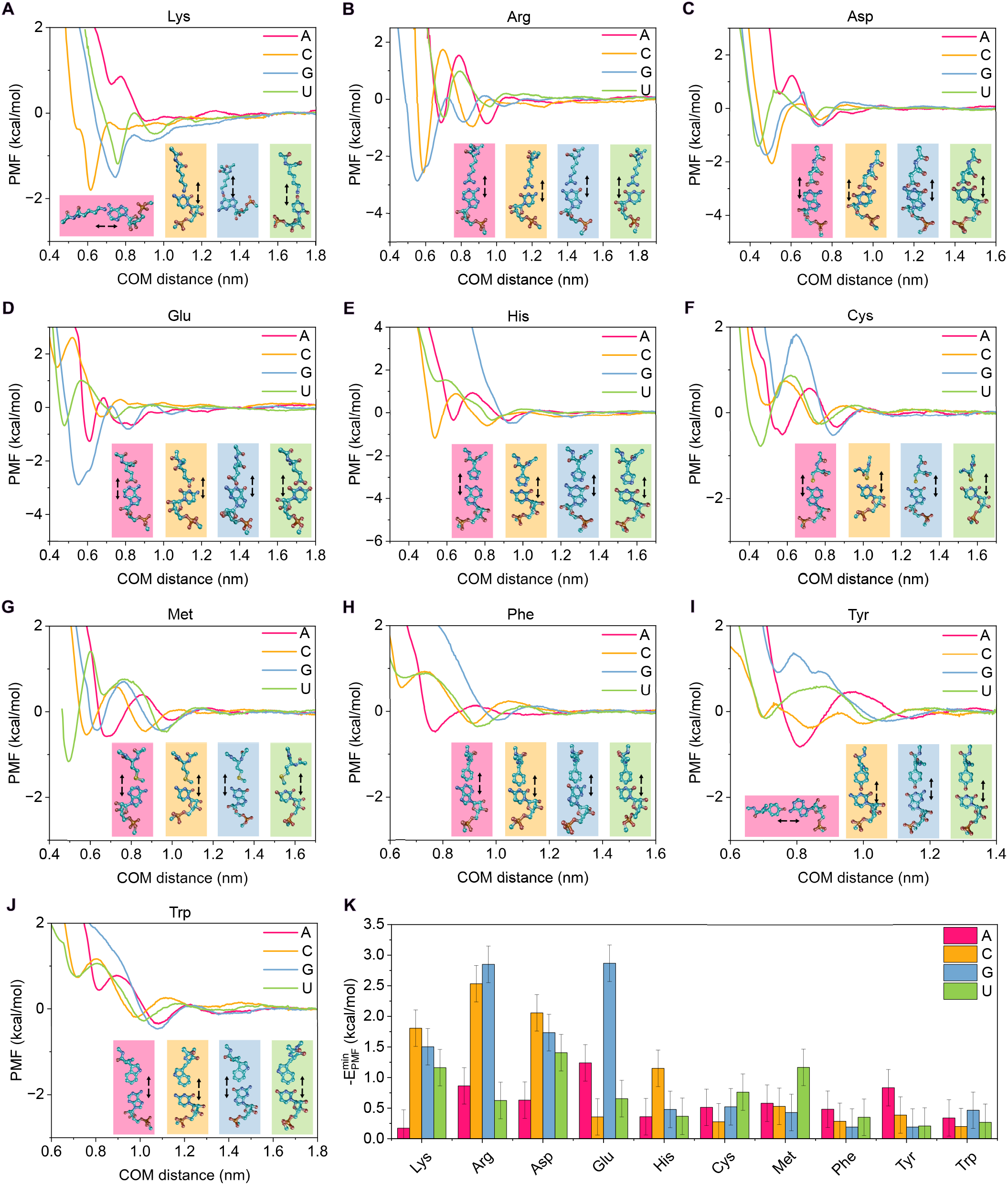
PMF profiles along the COM distance between the base of the nucleotide and the side chain of the amino acid obtained with AMBER03ws for the interactions between the amino acids lysine (**A**), arginine (**B**), aspartic acid (**C**), glutamic acid (**D**), histidine (**E**), cysteine (**F**), methionine (**G**), phenylalanine (**H**), tyrosine (**I**), and tryptophan (**J**) with the nucleotides adenine (pink curves), cytosine (yellow curve), guanine (blue curve), and uracil (green curve). The amino acids were oriented with their side chains facing the nitrogenous bases of the nucleotides, as shown next to each interaction profile including arrows to indicate the dissociation direction. **K** Absolute minimum PMF values of each nucleotide–amino acid pair from the interaction profiles shown in (**A**–**J**).

**FIG. 6.**
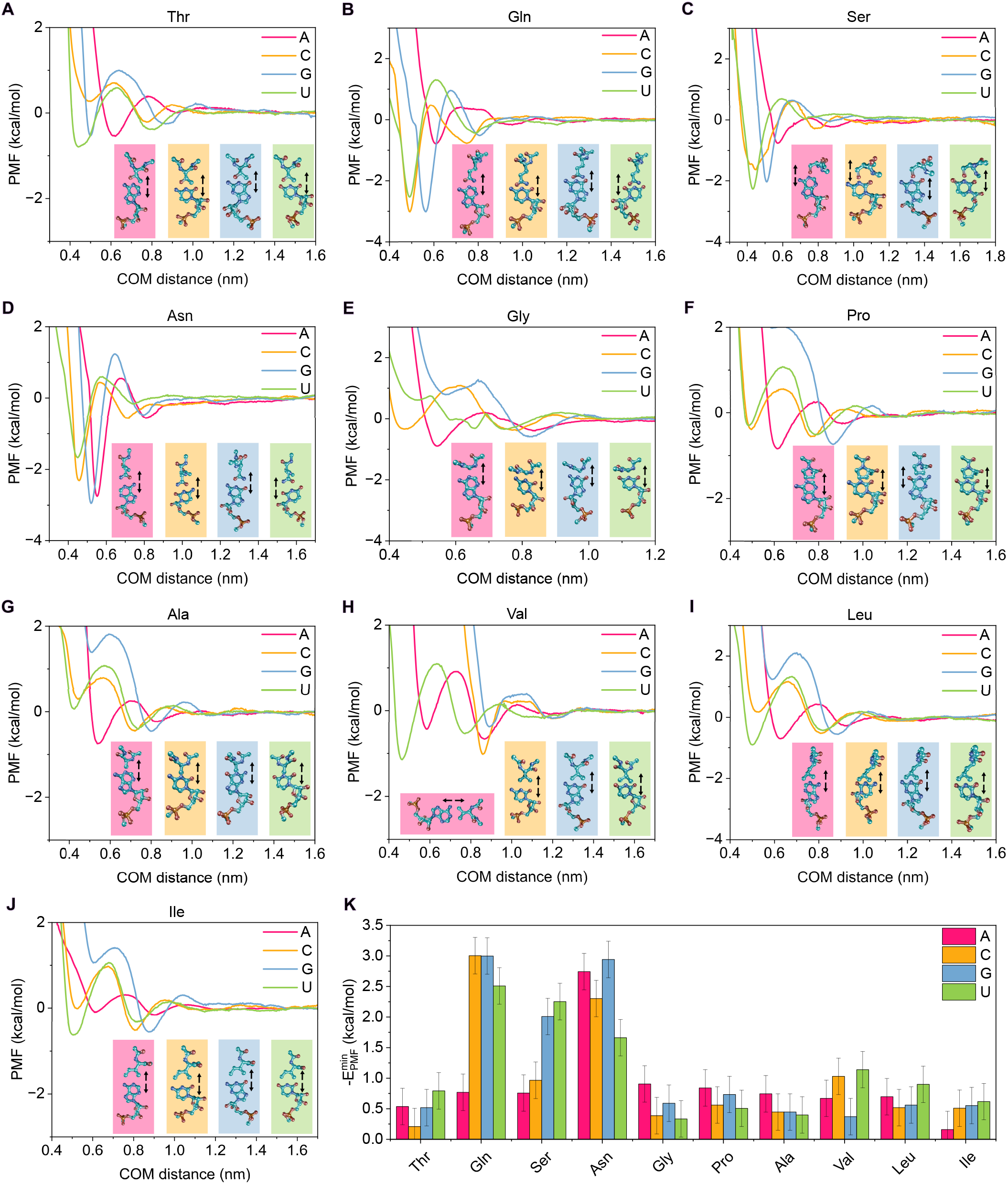
PMF profiles as a function of the COM distance between the base of the nucleotide and the side chain of the amino acid obtained with AMBER03ws for the interactions between the amino acids threonine (**A**), glutamine (**B**), serine (**C**), asparagine (**D**), glycine (**E**), proline (**F**), alanine (**G**), valine (**H**), leucine (**I**), and isoleucine (**J**) with the nucleotides adenine (pink curves), cytosine (yellow curve), guanine (blue curve), and uracil (green curve). The amino acids were oriented with their side chains facing the nitrogenous bases of the nucleotides, as shown next to each interaction profile including arrows to indicate the dissociation direction. **K** Absolute minimum PMF values of each nucleotide–amino acid pair from the interaction profiles shown in (**A**–**J**).

Taking a closer look at the positively charged amino acids (*i*.*e*., Arg and Lys; Figs. 5A–B,K), we observed that both arginine and lysine exhibited a strong affinity for C and G, whereas its binding to A and U was significantly weaker (≤1 kcal/mol). Moreover, we find that Arg interacts more strongly than Lys with all nucleotides, except uridine. This result is consistent with the expected role of arginine as a sticker, in contrast to lysine^26,112–115^. For example, arginine-rich regions are fundamental for the recruitment of RNA strands in FUS condensates^28,49,71^. Indeed, the demonstrated ability of Arg to form persistent cation–*π* and electrostatic interactions clearly explains its strong binding to nucleotides^58,112^. We attribute the stronger attraction between Arg and G/C to the greater polarizability of these bases compared to A and U. Interestingly, the interaction of histidine—with an effective charge of approximately +0.5 due to its partial protonation of the imidazole side chain (Fig. 5)—is negligible compared to arginine and lysine, with only cytosine showing moderate affinity. Moreover, negatively charged amino acids (Asp and Glu; Figs. 5C–D,K) also exhibited significant binding to certain nucleotides through its base. However, the negative charge of the nucleotide phosphate group can hinder direct interactions between these residues and the nucleotides when the binding does not occur through the base. We ascribe the attractive weak interactions to charge screening by solvent, ions and the exposure of the nucleotide base in this configuration, which likely reduces the electrostatic contrast between the two groups.

Aromatic residues (Tyr, Phe, and Trp) are typically expected to display high affinity toward RNA strands through *π*–*π* interactions^19,58,61^. However, our results reveal negligible binding (Figs. 5H–K), potentially due to the imposed orientation, which hinders *π*–*π* stacking (we will later discussed favorable binding modes which promote *π*–*π* interactions). In contrast, polar amino acids (Gln, Ser, Asn; Figs. 6B–D,K) exhibited significant binding energies (∼ 2–3 kcal/mol). For instance, the amide group in Asn can form hydrogen bonds with all nucleotides, as demonstrated by our calculations (Fig. 6D,K). Similarly, the amide group in Gln promotes persistent interactions with C, G, and U, whereas interactions with A are more limited due to the absence of carbonyl groups in its structure. Analogously, Ser can form hydrogen bonds between its hydroxyl group and the nucleobase. Interestingly, our results show that Thr interacts only weakly with the nucleotides (*E* ≲ 1 kcal/mol; Fig. 6A,K) despite its polar nature. This is consistent with the simpler chemical structure of threonine, which might render its side chain ineffective in stabilizing base interactions.

The remaining amino acids not discussed above (Cys, Met, Gly, Pro, Ala, Val, Leu, and Ile; Figs. 5F,G,K and 6C,E–K) exhibited negligible interactions (*E* ≲ 1 kcal/mol), thus contributing little to protein–RNA interactions that stabilize phase-separation. From the nucleotide perspective, cytosine (yellow) and guanine (blue) appear to be the most relevant residues in protein–RNA interactions, showing binding affinities of up to ∼3 kcal/mol with several amino acids (e.g., Arg, Glu, Gln). Uridine (green) can also interact significantly with representative amino acids, although the binding strengths are generally lower than those observed for C and G. In contrast, adenosine (red) displays only weak interactions, primarily its strongest binding occurs with Asn (Figs. 6K) and Glu (Figs. 5K), while the remaining amino acids exhibited negligible affinity toward it. Interestingly, polyA is among the most prevalent RNA strands in stress granules, where it recruits RNA-binding proteins such as TDP-43, FUS, or G3BP1^25,116–119^. Therefore, the main interaction mode of adenosine with amino acids is likely modulated through alternative mechanisms (i.e., non-specific electrostatic interactions through the phosphate group). To further analyze nucleotide–amino acid interactions involving the nucleotide base, we computed the integral of the negative part of the PMF curves (Fig. S6 in the SM). The overall trend observed for the PMF minima is preserved in the integral analysis. Notably, the most significant differences arise for polar residues (Gln, Ser, Asn), whose effective interactions appear weaker than in Figs. 5 and 6. This reduction can be attributed to the shape of the PMF curve at intermediate distances and the narrower width of the potential well at the global minimum.

Furthermore, we focused on alternative specific nucleotide–amino acid interactions that are particularly relevant to biomolecular condensation. In particular, we explored: (i) the interactions between charged amino acids (Arg, Lys, Glu, Asp) and the phosphate group of the nucleotide (Fig. 7B); (ii) the binding of aromatic residues (Tyr, Phe) to the nucleobase through *π*–*π* stacking (Fig. 7A); and (iii) the interaction between tyrosine and the ribose moiety of cytosine and uridine (Fig. 7C). The corresponding PMF curves are shown in Fig. S7 of the SM, together with representative snapshots of the configurations employed. Our calculations for *π*–*π* stacking interactions between the aromatic rings of Tyr and Phe and the nitrogenous bases of the nucleotides (Fig. 7A) revealed strong binding energies (5–6 kcal/mol), which are similar across all nucleotide–amino acid pairs. Remarkably, these interaction energies are comparable to those associated with CG base pairing (Fig. 1), underscoring the recognized role of aromatic contacts in the formation and stabilization of RNA–protein condensates^28,58,120^ and consistent with previous computational PMF calculations^61^.

**FIG. 7.**
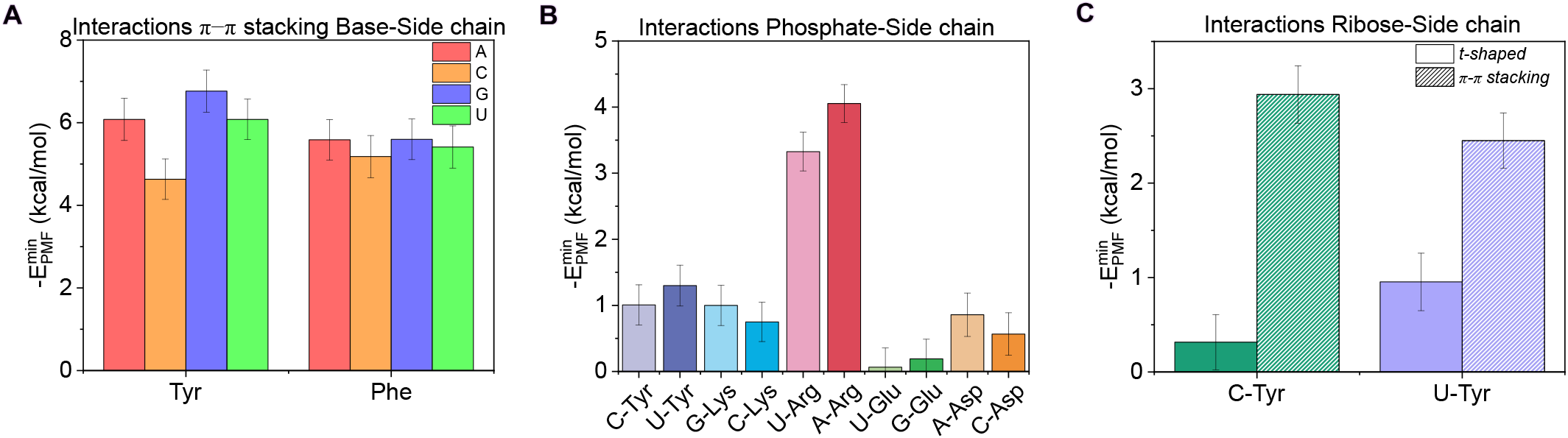
(**A**) PMF values from *π*–*π* stacking interactions between the bases of the nucleotides and the side chains of the aromatic amino acids tyrosine and phenylalanine with adenine (red bars), cytosine (orange bars), guanine (blue bars), and uracil (green bars). (**B**) Interaction PMF minima of selected nucleotide–amino acid pairs via the phosphate groups of the nucleotides and the side chains of the amino acids. (**C**) Interaction PMF minima between ribose moieties of the nucleotides and the side chains of the amino acids, oriented in t-shaped conformations (solid bars) and *π*–*π* stacking conformations (striped bars), for tyrosine interacting with cytosine (green bars) and uracil (purple bars). All the calculations were performed with AMBER03ws at 300 K, 1 bar, and 150 mM NaCl.

To evaluate the electrostatic interactions (Fig. 7B), we employed different representative nucleotides for each of the charged amino acids (Arg, Lys, Glu, Asp), and also included the aromatic residue Tyr as a cross-check. In principle, the specific identity of the nucleotide should not dramatically affect the interaction between the phosphate group and the amino acid side chain; therefore, only two nucleotides were used for the various amino acids. Our results showed that the interaction with arginine (red) is the strongest, reaching approximately 3.5–4.0 kcal/mol. Interestingly, the computed binding PMF profiles between the phosphate group and the amino acid side chain suggests unfavorable interactions in several cases (U–Glu, G–Glu, and C–Asp). Lysine, despite its positively charged side chain, only showed a moderately attractive interaction for the phosphate group compared to arginine. This can be attributed to the entropic penalty associated with forming a stable contact: lysine must first displace tightly bound water molecules, and the energy gained upon forming the Lys–phosphate bridge might not sufficiently compensate for this cost to represent a strong energetic gain. These findings reinforce previous experimental observations highlighting the dominant role of arginine, as opposed to lysine, in driving RNA–protein phaseseparation^26,112,113,121^. Finally, our results confirm that the interaction between the phosphate group and amino acids is largely independent of the specific nucleotide considered.

Finally, we calculated the interaction between tyrosine and the ribose moiety of cytosine and uridine, considering two distinct conformations (see Fig. S7 for representative snapshots): one in which both rings are stacked (referred to as *π*–*π* stacking), and another where the tyrosine side chain is oriented perpendicular to the ribose plane (t-shaped; Fig. S7D). For both nucleotide–Tyr pairs, we observed that the *π*–*π* stacking interactions were significantly stronger than the t-shaped ones, although these *π*–*π* interactions were weaker than those between the aromatic ring of Tyr and the nitrogenous bases of C and U (Fig. 7A). Importantly, while the t-shaped conformation yielded negligible binding compared to interactions typically considered significant (*E* ≳ 2 kcal/mol), the *π*–*π* stacking between Tyr and the ribose displayed comparable strength to amino acid–amino acid interactions such as Tyr-Tyr interactions, thereby notably contributing to the stabilization of RNA–protein condensates.

## CONCLUSIONS

In this work, we presented an extensive analysis of nucleotide and amino acid interactions relevant to the formation of biomolecular condensates. Specifically, we investigated several configurations of nucleotide–nucleotide, amino acid–amino acid, and nucleotide–amino acid pairs by applying potential-of-mean-force calculations in atomistic MD simulations. We examined different interaction modes of nucleotide–nucleotide pairs by computing PMFs for canonical Watson–Crick pairs (CG and AU) as well as for the GU wobble pair, using both AMBER03ws/TIP4P2005 and CHARMM36/TIP4P2005 force-field combinations (Fig. 1). Our results showed excellent agreement between the two force fields and reproduced the expected energetic hierarchy according to the hydrogen bonds number that each canonical base-pair can establish (CG *>* AU *>* GU). Furthermore, the analysis of non-canonical base pairs revealed that CC interactions are the most energetically favorable among them, while other non-canonical pairs exhibited variable but generally lower stability compared to the canonical ones (Fig. 1). A geometric analysis based on the C1’–C1’ distance further supported these energetic trends (Fig. S1), in agreement with experimental observations^90–92^.

We also analyzed alternative binding modes of the nucleotides through their ribose and phosphate groups, considering both *cis* and *trans* conformations to assess the effect of nucleotide orientation (Fig. 2A-D). Ribose–ribose interactions exhibited modest attractive energies, with *trans* orientations generally being more favorable due to reduced electrostatic repulsion. Similarly, phosphate–phosphate interactions displayed PMF profiles with shallow energy wells, reflecting the significant electrostatic repulsion between negatively charged groups. Remarkably, *cis* configurations were more stable than *trans* orientations, and among them, pairs involving guanine exhibited the most favorable (though still weak) binding affinity. RNA tertiary interactions were further characterized by computing the energetic landscape of a G-quadruplex structure (Fig. 2E). Our results demonstrated strong cooperative interactions among the RNA strands, yielding strong binding strengths comparable to those reported for protein cross-*β*-sheets^50,66^. Crucially, this high stability arises from the formation of multiple Hoogsteen-type hydrogen bonds between guanine units. Finally, we examined the effect of salt concentration on the various nucleotide interaction modes (Fig. 3). Increasing salt concentration leads to a moderate reduction in the binding between bases and riboses, and more notably, a pronounced weakening of phosphate–phosphate repulsive interactions due to screening of short-range electrostatic forces.

We further characterized residue–residue interactions by extending our PMF calculations to amino acids. First, we benchmarked amino acid pair interactions using AMBER03ws against the a99SB-*disp* force field, finding near-quantitative agreement (Fig. 4). We then estimated the interactions between all 20 amino acids and the four RNA nucleotides by orienting the nucleobase toward the amino acid side chain (Figs. 5 and 6). Only a subset of amino acids (Lys, Arg, Asp, Glu, Gln, Ser, and Asn) established significantly attractive interactions with most nucleotides, primarily driven by hydrogen bonding and cation–*π* interactions. To complement this analysis, we examined specific biologically relevant interactions, including *π*–*π* stacking, phosphate–side chain contacts, and ribose–side chain interactions (Fig. 7). We identified *π*–*π* stacking between Tyr or Phe and nucleobases as a strong and relevant binding mode involved in phase separation^58,61^. Moreover, electrostatic interactions with the phosphate group were particularly significant for arginine, in agreement with experimental observations of protein–RNA condensates^26,113^. Finally, ribose–*π* stacking emerges as a major potential RNA–protein interaction in biomolecular condensates. Taken together, our results revealed specific affinity patterns between nucleotides and amino acids, providing insight into how residue-level interactions influence molecular organization at larger scales in biomolecular condensates.

## Data availability

We provide the relevant data in the repository ([GitHub link for the repository](https://github.com/Reshiiiii/PMF_RNA_Data_Scripts)) to facilitate reproducibility of our results. In the repository we also give the necessary code to run the simulations.

## ACKNOWLEDGEMENTS

A. F. acknowledges funding from the Ramon y Cajal fellowship (RYC2021-030937-I) and the project PID2022-136919NA-C33 from the Spanish MICIN. E.P. acknowledges funding from European Social Fund Plus and the project PID2022-136919NA-C33 from the Spanish MICIN. I. S.-B. acknowledges funding from the Derek Brewer scholarship of Emmanuel College and EPSRC Doctoral Training Programme studentship, number EP/T517847/1, Ramon y Cajal fellowship (awarded to J.R.E.), as well as the UKRI EPSRC under the UK Government’s guarantee scheme (EP/Z002028/1), following successful evaluation by the ERC (Consolidator Grant awarded to R.C.G.) under the European Union’s Horizon Europe research and innovation programme. A.R acknowledges funding from project PID2023-147156NB-I00 from the Spanish MICIN. R.C.-G. acknowledges funding from the European Research Council (ERC) under the European Union Horizon 2020 research and innovation programme (grant agreement 803326). A. T. is funded by European Research Council (ERC) under the European Union Horizon 2020 research and innovation programme (grant agreement 803326) and Ramon y Cajal fellowship (RYC2021-030937-I). J. R. E. also acknowledges funding from the Ramon y Cajal fellowship (RYC2021-030937-I), the Spanish National Agency for Research (PID2022-136919NA-C33), and the European Research Council (ERC) under the European Union’s Horizon Europe research and innovation program (grant agreement no. 101160499). This work has been performed using resources provided by the Cambridge Tier-2 system operated by the University of Cambridge Research Computing Service (http://www.hpc.cam.ac.uk) funded by EPSRC Tier-2 capital grant EP/P020259/1. The authors also thankfully acknowledge RES computational resources provided by MareNostrum 5 through the activities FI-2024-3-0001 and FI-2025-1-0008.

