## Supplementary Material for "Determination of nucleotide-nucleotide and nucleotide-amino acid binding interactions from all-atom potential-of-mean-force calculations"

Ignacio Sanchez-Burgos

*Yusuf Hamied Department of Chemistry, University of Cambridge,*

*Lensfield Road, Cambridge CB2 1EW, UK.*

Rosana Collepardo-Guevara

*Yusuf Hamied Department of Chemistry, University of Cambridge,*

*Lensfield Road, Cambridge CB2 1EW, UK. and*

*Department of Genetics, University of Cambridge,*

*Downing Street, Cambridge CB2 3EH, UK.*

Andrés R. Tejedor[*]

*Yusuf Hamied Department of Chemistry, University of Cambridge,*

*Lensfield Road, Cambridge CB2 1EW, UK. and*

*Department of Physical Chemistry, Universidad Complutense de Madrid,*

*Av. Complutense s/n, Madrid 28040,*

*Spain.* [†]*These authors contributed equally*

Jorge R. Espinosa[†]

*Department of Physical Chemistry, Universidad Complutense de Madrid,*

*Av. Complutense s/n, Madrid 28040,*

*Spain.* [†]*These authors contributed equally and*

*Yusuf Hamied Department of Chemistry, University of Cambridge,*

*Lensfield Road, Cambridge CB2 1EW, UK.*

### SUPPLEMENTAL FIGURES

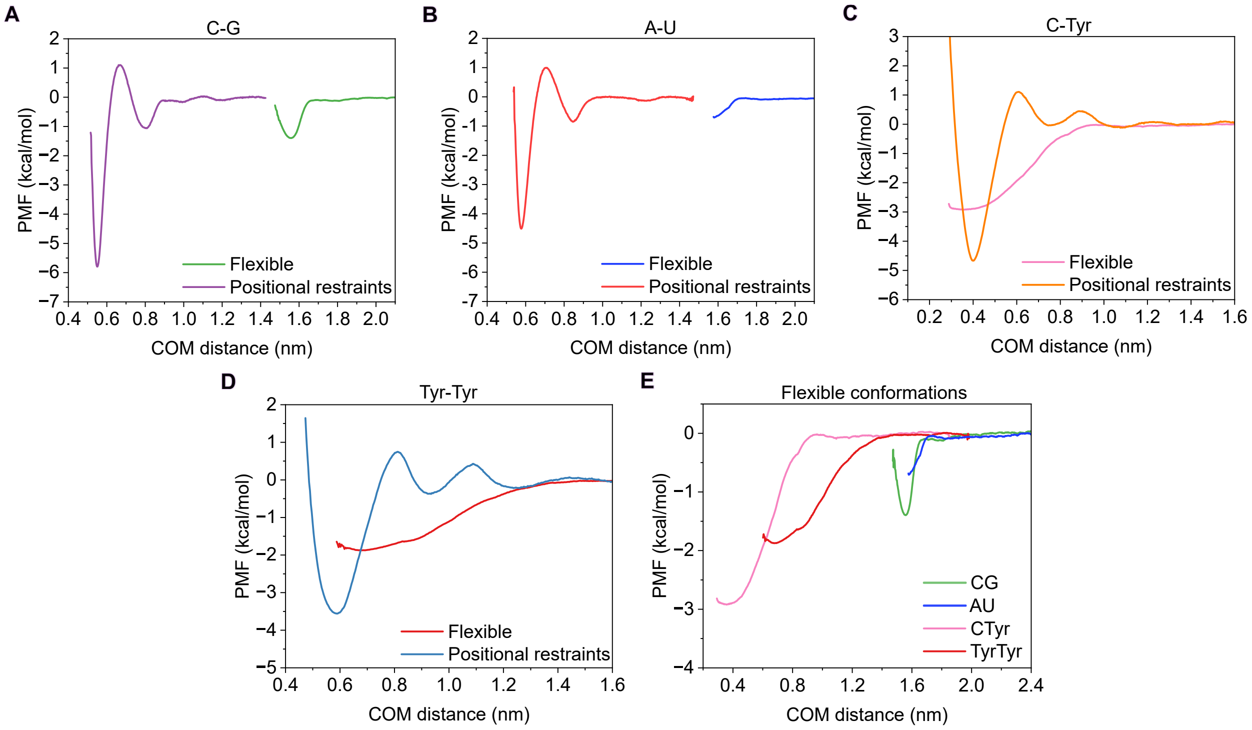


**FIG. S1:** Difference of atomistic potential of mean force (PMF) dissociation curve along the COM distance between the bases or side chain of the corresponding nucleotides or amino acids, respectively, at room conditions and physiological NaCl concentration (150 mM) in explicit solvent an ions with positional restraints in heavy atoms or only in the part of the molecule that is not confronted for the cytosine-guanine pair **(A)**, adenine-uracil pair **(B)** and cytosine-tyrosine pair **(C)** with the amber03ws force field and tyrosine-tyrosine pair **(D)** with the a99sb-*disp* force field. **(E)** Curve of PMFs for the flexible pairings. The center of mass (COM) and the positional restraints of these flexible conformations have been defined in the part of the molecule that is not confronted.


*

†

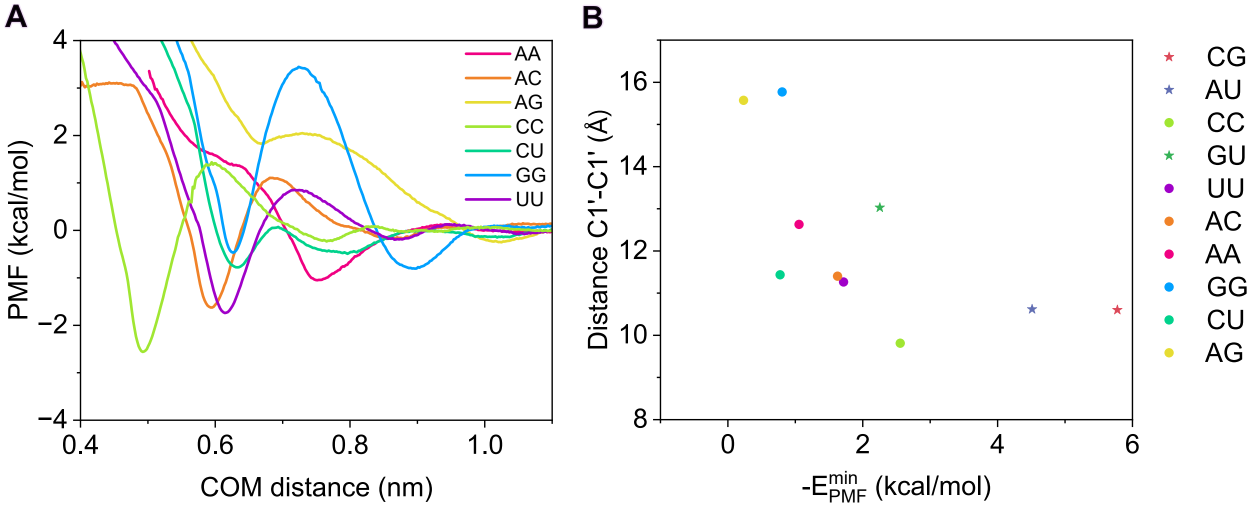


**FIG. S2:** (**A**) Atomistic potential of mean force (PMF) dissociation curve of the different nucleotides non-canonical pairs studied as a function of the center-of-mass (COM) distance between the bases of the corresponding pair using AMBER03ws force field. PMF simulations have been conducted at room conditions and physiological NaCl concentration (150 mM) in explicit solvent and ions. Nucleotides have been confronted with their bases. (**B**) Distance C1'-C1' vs. Minimum energy of the PMF curve for the different pairs of nucleotides.

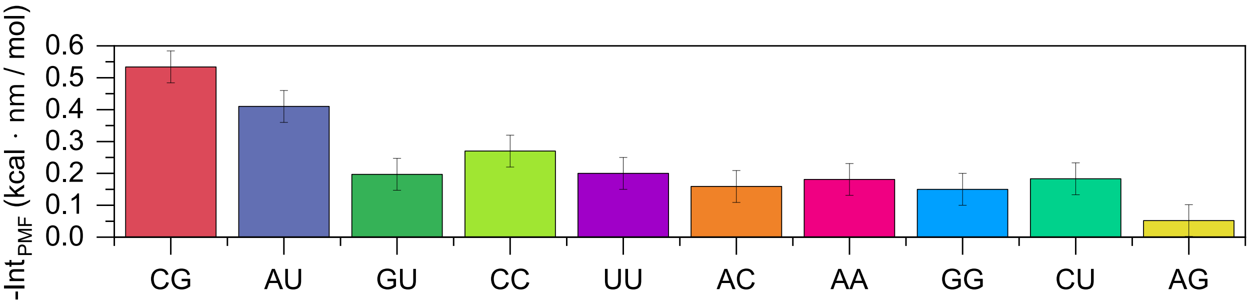


**FIG. S3:** Integrals of the attractive part of the corresponding PMF curves for canonical and non-canonical nucleotide pairs. All calculations were performed at 300 K and 150 mM NaCl with the AMBER03ws force field.

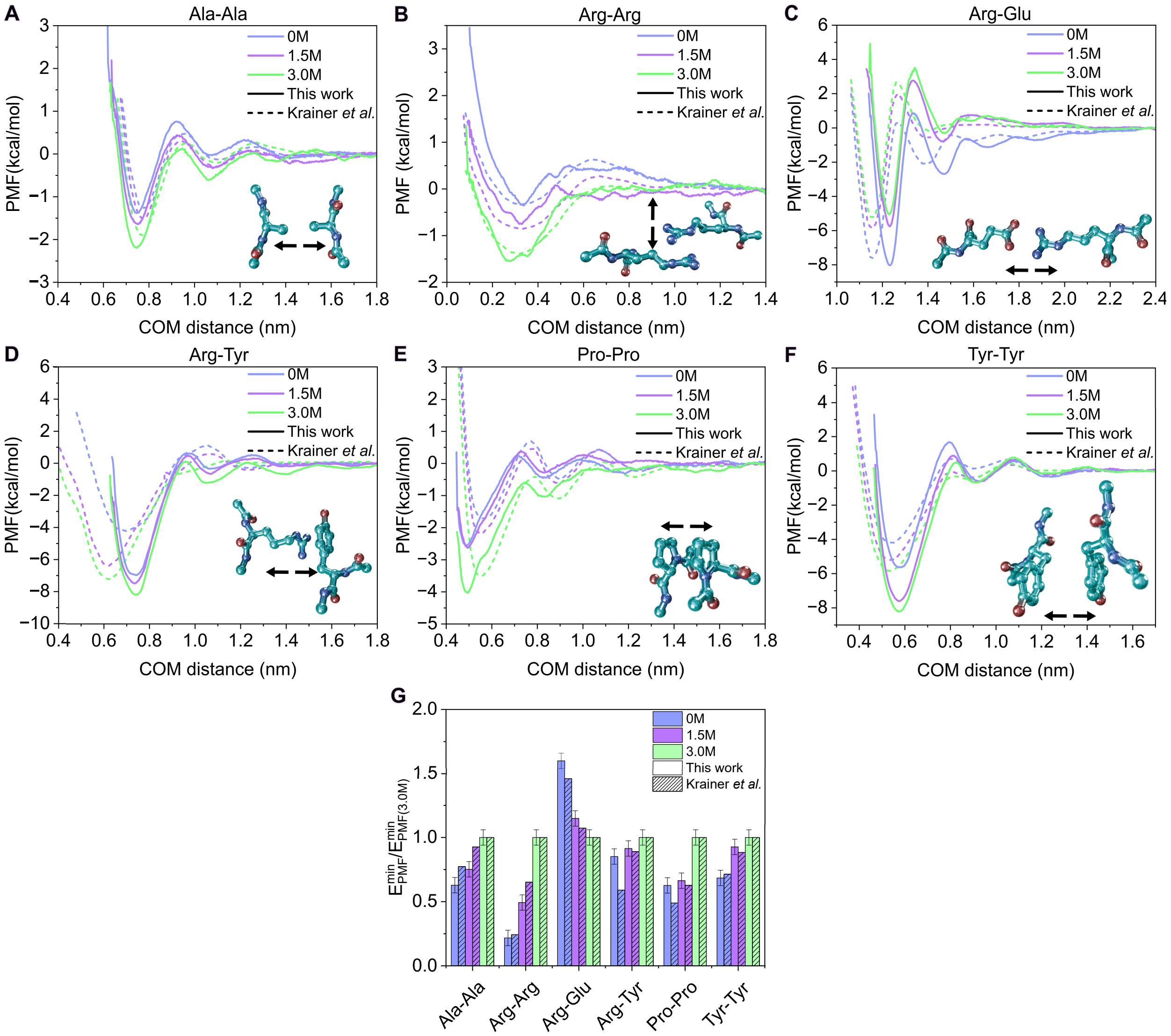


**FIG. S4:** Effect of salt concentration of NaCl 0 M (indigo), 1.5 M (violet) and 3 M (light green) on the atomistic potential of mean force (PMF) along the COM distance between the side chain of the corresponding amino acid pairs in explicit solvent and NaCl ions at room temperature as a function of the center-of-mass (COM) distance using AMBER03ws force field. The selected amino acids are alanine-alanine (**A**), arginine-arginine (**B**), arginine-glutamic (**C**), arginine-tyrosine (**D**), proline-proline (**E**) and tyrosine-tyrosine (**F**). The data from this work have been plotted in Figures **A-F** with a solid line and the results of Ref. [1] in a dotted line. In Figs **A-F** as an inset we have represented the different configurations of these pairs including arrows to indicate the dissociation direction. The minimum energy of the PMF normalized against the value of that pair at 3.0 M has been plotted for the different pairs in **G**. In this Figure the results of this work have been represented with solid bars, and the results of Ref. [1] with dashed bars.

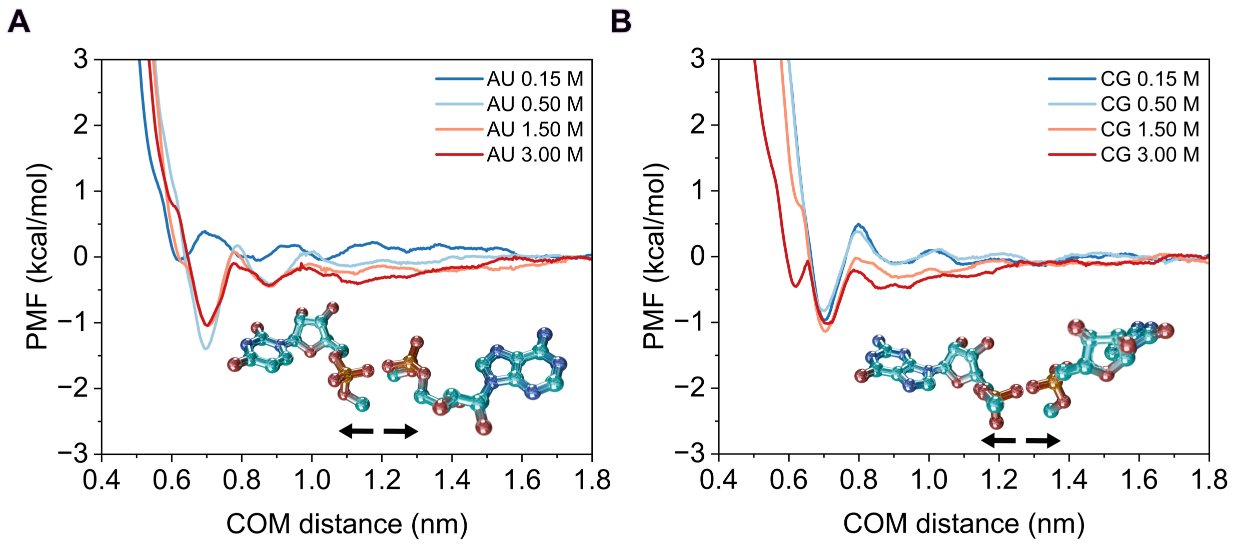


**FIG. S5:** PMF along the COM distance between the phosphate group of the corresponding nucleotides at different concentrations with AMBER03ws for the canonical nucleotide pairs AU (**A**) and CG (**B**), oriented via their phosphate groups. As an inset we have represented the different configurations of these pairs including arrows to indicate the dissociation direction.

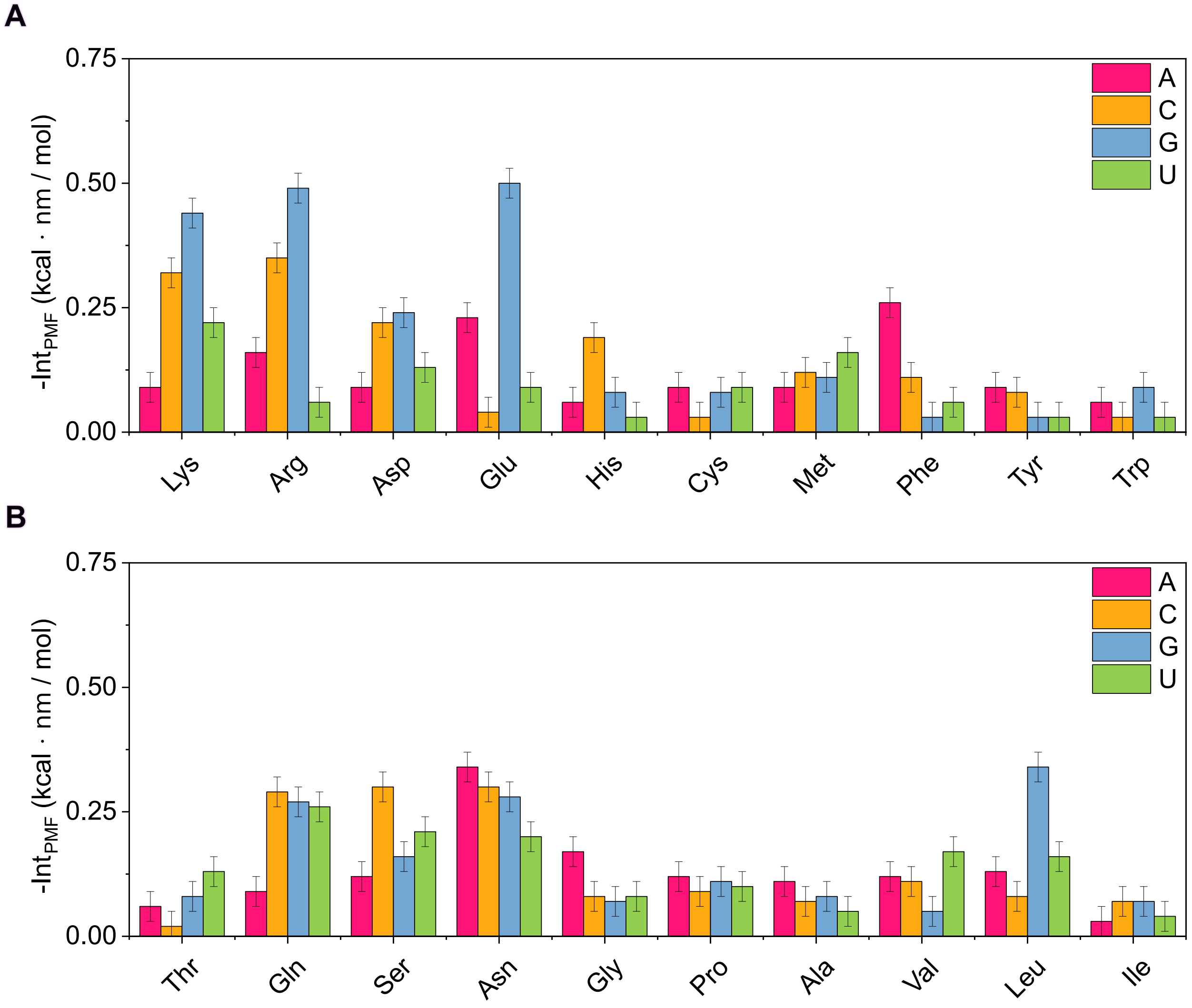


**FIG. S6:** Integrals of the attractive part of the corresponding PMF curves of each nucleotide–amino acid pair are shown. Aminoacids (**A**) Lysine, arginine, aspartic acid, glutamic acid, histidine, cysteine, methionine, phenylalanine, tyrosine, and tryptophan, (**B**) threonine, glutamine, serine, asparagine, glycine, proline, alanine, valine, leucine, and isoleucine with the nucleotides adenine (pink bars), cytosine (yellow bars), guanine (blue bars), and uracil (green bars).

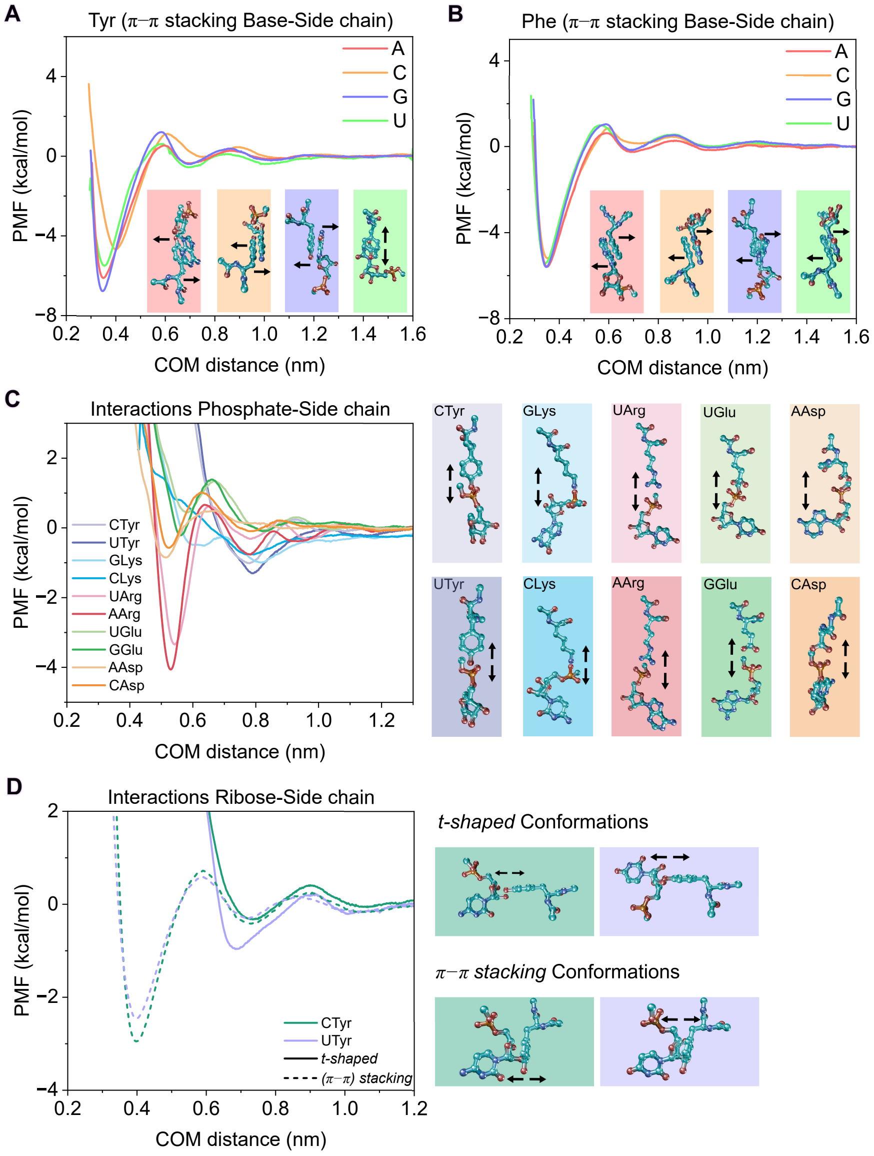


**FIG. S7:** PMF dissociation curve at 300K and 150 mM of NaCl in explicit solvent an ions using AMBER03ws force field between selected nucleotides and amino acids. Interaction via $\pi$-$\pi$ stacking of the nucleotide's base and the amino acid's side chain for the different nucleotides with tyrosine (**A**) and phenilalanine (**B**). (**C**) PMF curve of the confronted nucleotide's phosphate and the amino acid's side chain between different nucleotides and amino acids. (**D**) PMF curve of the confronted nucleotide's ribose and the amino acid's side chain between the tyrosine with cytosine and uracil. In solid curves we plotted the t-shaped conformations and in dotted curve the $\pi$-$\pi$ stacking conformations. In Figures **C** and **D** on the right side we show the different configurations of these pairs including arrows to indicate the dissociation coordinate.
